# Hyperglycemia increases the severity of *Staphylococcus aureus* osteomyelitis and influences bacterial genes required for survival in bone

**DOI:** 10.1101/2022.11.30.518635

**Authors:** Casey E. Butrico, Nathan B. Klopfenstein, Erin R. Green, Joshua R. Johnson, Sun H. Peck, Carolyn B. Ibberson, C. Henrique Serezani, James E. Cassat

## Abstract

Hyperglycemia, or elevated blood glucose, renders individuals more prone to developing severe *Staphylococcus aureus* infections. *S. aureus* is the most common etiological agent of musculoskeletal infection, which is a common manifestation of disease in hyperglycemic patients. However, the mechanisms by which *S. aureus* causes severe musculoskeletal infection during hyperglycemia are incompletely characterized. To examine the influence of hyperglycemia on *S. aureus* virulence during invasive infection, we used a murine model of osteomyelitis and induced hyperglycemia with streptozotocin. We discovered that hyperglycemic mice exhibited increased bacterial burdens in bone and enhanced dissemination compared to control mice. Furthermore, infected hyperglycemic mice sustained increased bone destruction relative to euglycemic controls, suggesting that hyperglycemia exacerbates infection-associated bone loss. To identify genes contributing to *S. aureus* pathogenesis during osteomyelitis in hyperglycemic animals relative to euglycemic controls, we used transposon sequencing (TnSeq). We identified 71 genes uniquely essential for *S. aureus* survival in osteomyelitis in hyperglycemic mice and another 61 mutants with compromised fitness. Among the genes essential for *S. aureus* survival in hyperglycemic mice was superoxide dismutase A (*sodA*), one of two *S. aureus* superoxide dismutases involved in detoxifying reactive oxygen species (ROS). We determined that a *sodA* mutant exhibits attenuated growth *in vitro* in high glucose and *in vivo* during osteomyelitis in hyperglycemic mice. SodA therefore serves an important role during growth in high glucose and promotes *S. aureus* survival in bone. Collectively, these studies demonstrate that hyperglycemia increases the severity of osteomyelitis and identify genes contributing to *S. aureus* survival during hyperglycemic infection.

## INTRODUCTION

Hyperglycemia, or elevated blood glucose, can result from chronic metabolic conditions as well as acute stress and reflects the inability of the body to produce or effectively use insulin. Hyperglycemia leads to a variety of physiological and immune regulatory alterations.^1–6^ Dysregulated cytokine response, altered chemotaxis, and decreased function of innate immune cells contribute to a hyperinflammatory environment and influence the pathophysiology of bacterial infections.^1,4,7^ Hyperglycemia also increases oxidative stress in tissues via production of reactive oxygen species (ROS), which are generated as a consequence of electron transport chain dysfunction during mitochondrial respiration and glucose metabolism.^8^ The combination of reduced immune responses and hyperinflammation leads to a to greater incidence of infection in individuals with hyperglycemia.^9^ *S. aureus* is a particularly common etiologic agent of severe infections in patients with hyperglycemia. Infection of bone, or osteomyelitis, is one of the most frequent manifestations of staphylococcal infection in these patients.^10^ *S. aureus* osteomyelitis is particularly difficult to treat due to widespread antibiotic resistance, antibiotic tolerance, and the induction of bone destruction that can limit antibiotic delivery to the infectious focus.^11,12^ Additionally, individuals with chronic hyperglycemia have greater bone porosity, higher rates of osteoporosis, and enhanced risk of bone fractures, further complicating treatment.^13,14^ However, how hyperglycemia alters osteomyelitis pathogenesis or bacterial adaptation to the host microenvironment during osteomyelitis is not fully understood.

*S. aureus* has a remarkable ability to infect a variety of tissues, which can be attributed to its metabolic flexibility and the capacity to produce an arsenal of virulence factors.^15–18^ Multiple transcriptional regulators allow *S. aureus* to modulate its virulence in response to environmental cues, such as glucose abundance.^17^ However, the virulence and metabolic mechanisms by which *S. aureus* adapts to the altered host environment during osteomyelitis in hyperglycemic mice are not well understood. We hypothesized that the alterations in glucose availability and changes in host physiology during hyperglycemia require *S. aureus* to use distinct genes to survive and induce severe disease in the context of osteomyelitis.

To investigate the requirements for *S. aureus* survival and virulence *in vivo* during hyperglycemia, we subjected mice to acute and chronic hyperglycemia and then induced osteomyelitis using a post-traumatic model. Hyperglycemic infected mice exhibited more severe infection as measured by CFU burden and bone destruction. We used transposon sequencing (TnSeq) to identify *S. aureus* genes required for infection in hyperglycemic osteomyelitis. Based on the TnSeq analysis, we further studied the influence of superoxide dismutase A (SodA) on disease pathogenesis due to the known role of SodA in detoxifying ROS.^19^ SodA was found to be important for *S. aureus* survival in hyperglycemic mice. To study the mechanistic basis of this finding, we analyzed the growth of a *sodA* mutant *in vitro* in high glucose and identified that the *sodA* mutant exhibited a growth defect in high glucose. Collectively, this study uncovers mechanisms of increased virulence during *S. aureus* osteomyelitis in hyperglycemic mice.

## RESULTS

### Acute hyperglycemia increases *S. aureus* burdens during osteomyelitis

To investigate the impact of hyperglycemia on *S. aureus* virulence during osteomyelitis, we first induced hyperglycemia in male wildtype (WT) mice by treating with streptozotocin (STZ).^7,20^ STZ induces hyperglycemia through cytotoxic effects on insulin-producing beta cells. We then subjected STZ- or vehicle-treated mice to osteomyelitis and determined *S. aureus* burdens following infection.^18^ Because the induction of hyperglycemia following STZ treatment is variable, we measured blood glucose levels in mice 10 days after STZ treatment, at the time of infection (day 0) and at the conclusion of the experiment (day 14).^21,22^ In this study, all mice treated with STZ were hyperglycemic (defined as blood glucose levels of > 250 mg/dl) at days 0 and 14 of infection (**Fig. 1A**). STZ-treated hyperglycemic mice and euglycemic vehicle-treated control mice were infected 10 days after the last STZ or vehicle treatment with 1×10^6^ CFU *S. aureus* USA300 lineage strain AH1263 (WT). To infect mice, we used a post-traumatic model of osteomyelitis, in which bacteria are inoculated directly into a cortical defect in the mid-femur.^18,23^ Acute hyperglycemia resulted in a significant increase in *S. aureus* burdens in the infected femurs compared to euglycemic vehicle-treated animals at day 14 post-infection (**Fig. 1B**). To assess the extent to which hyperglycemia alters *S. aureus* dissemination to other tissues, we also collected the contralateral femur, kidneys, liver, and heart. While dissemination to the contralateral femur did not significantly change, dissemination to the kidneys, liver, and heart significantly increased in acute hyperglycemic animals (**Fig. 1C-F**). Similar trends were observed with a lower inoculum of *S. aureus* (**Fig. S1**), although the only organs with significantly greater CFU in the infected hyperglycemic mice compared to euglycemic vehicle-treated were the infected femur and kidneys (**Fig. S1B and S1C**). These data suggest that acute hyperglycemia increases *S. aureus* survival in bone as well as dissemination to other organs during osteomyelitis.

**Figure 1.**
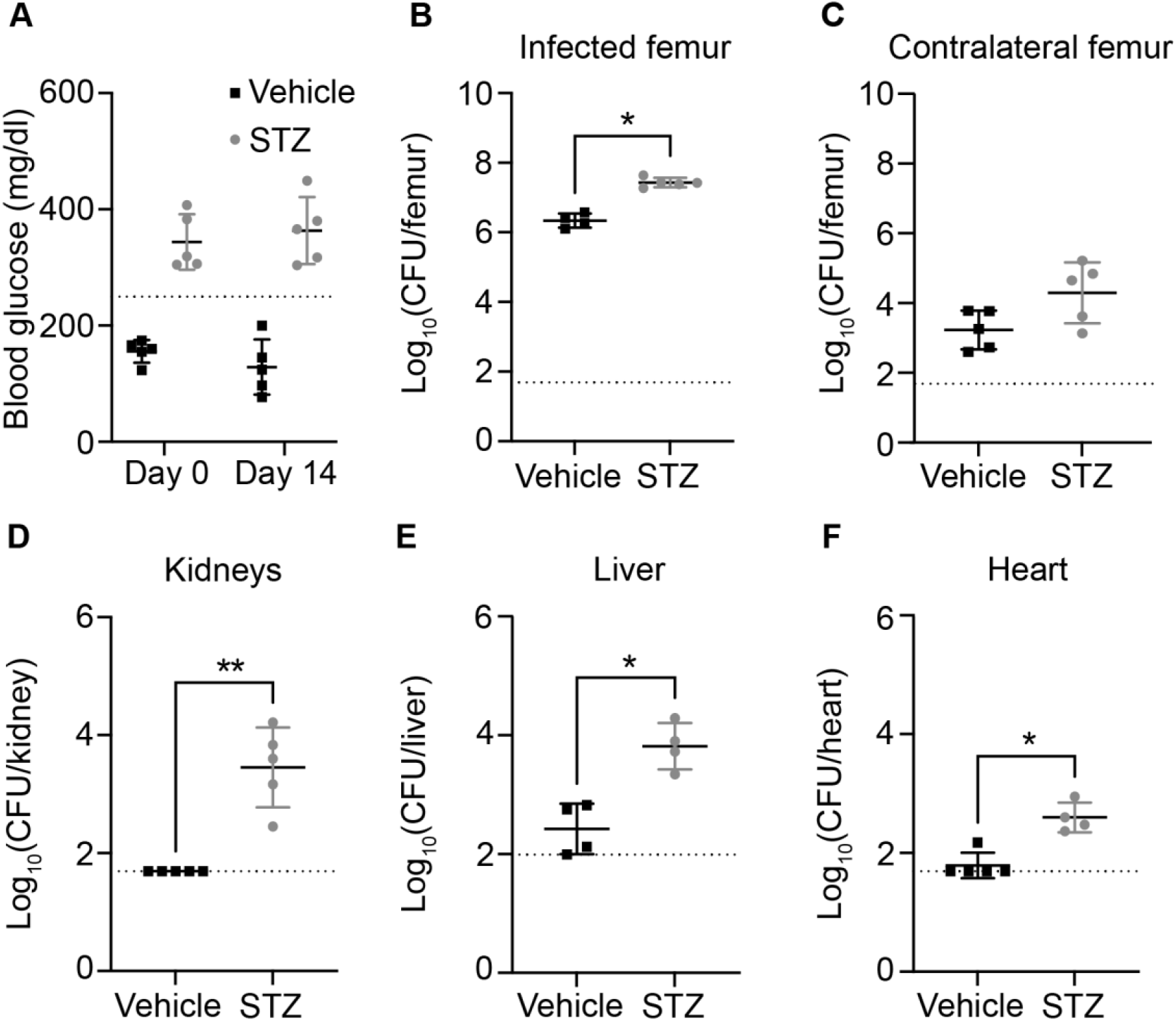
Acute hyperglycemia increases *S. aureus* burdens and dissemination during osteomyelitis. Eight-week old male mice were treated with sodium citrate (vehicle) or streptozotocin (STZ) intraperitoneally for 5 days. 10 days after the final injection, mice were infected with 1×10^6^ CFU of WT *S. aureus* via intraosseous injection. (A) Blood glucose concentration was quantified from a tail vein bleed immediately prior to inoculation (day 0) and the day of sacrifice (day 14). Dotted line indicates hyperglycemia threshold of 250 mg/dl. Mice were sacrificed at day 14 post-infection, and the bacterial burdens (CFU) were enumerated in (B) infected femur, (C) contralateral femur, (D) kidneys, (E) liver, and (F) heart. N = 5 mice per group. Dotted lines indicate limit of detection. Horizontal lines indicate mean, and error bars represent SD. Significance determined with Mann-Whitney test (B-F). *p<0.05, **p<0.01.

To confirm that changes in *S. aureus* growth and dissemination result from acute hyperglycemia induced from STZ treatment and not from off-target effects of the drug, we assessed bacterial burdens in three additional groups of mice: euglycemic vehicle-treated mice, STZ-treated mice that became hyperglycemic, and STZ-treated mice that remained euglycemic (**Fig. S2A**). We inoculated mice with 1×10^6^ CFU of *S. aureus* and measured bacterial burdens in the femur 14 days post-infection. STZ-treated hyperglycemic mice had greater *S. aureus* burdens in infected femurs compared to both euglycemic vehicle-treated and euglycemic STZ-treated mice (**Fig. S2B**). These data indicate that changes in bacterial burden in STZ-treated mice result from hyperglycemia.

### Chronic hyperglycemia increases *S. aureus* burden during osteomyelitis

Prior studies using STZ-induced hyperglycemia induced *S. aureus* infection 30 days after treatment.^1,24^ To determine if a more chronic state of hyperglycemia alters the pathogenesis of staphylococcal osteomyelitis, we treated mice with STZ for 5 days and then initiated osteomyelitis 30 days after the final STZ injection. Hyperglycemia was confirmed at the start (day 0) and end (day 14) points of the infection (**Fig. 2A**). As observed with the acute model of hyperglycemia, bacterial burdens were elevated in infected femurs from hyperglycemic mice compared to euglycemic vehicle-treated mice at day 14 post-infection (**Fig. 2B**). Dissemination to contralateral femurs did not increase in hyperglycemic infected mice (**Fig. 2C**). However, *S. aureus* burdens in the kidneys, liver, and heart of the hyperglycemic infected mice were increased compared to euglycemic mice at day 14 post-infection (**Fig. 2D-F**). Similar trends were observed using a ten-fold lower inoculum (**Fig. S3**). Taken together, these data suggest that both acute and chronic hyperglycemia result in increased *S. aureus* bacterial burdens during osteomyelitis.

**Figure 2.**
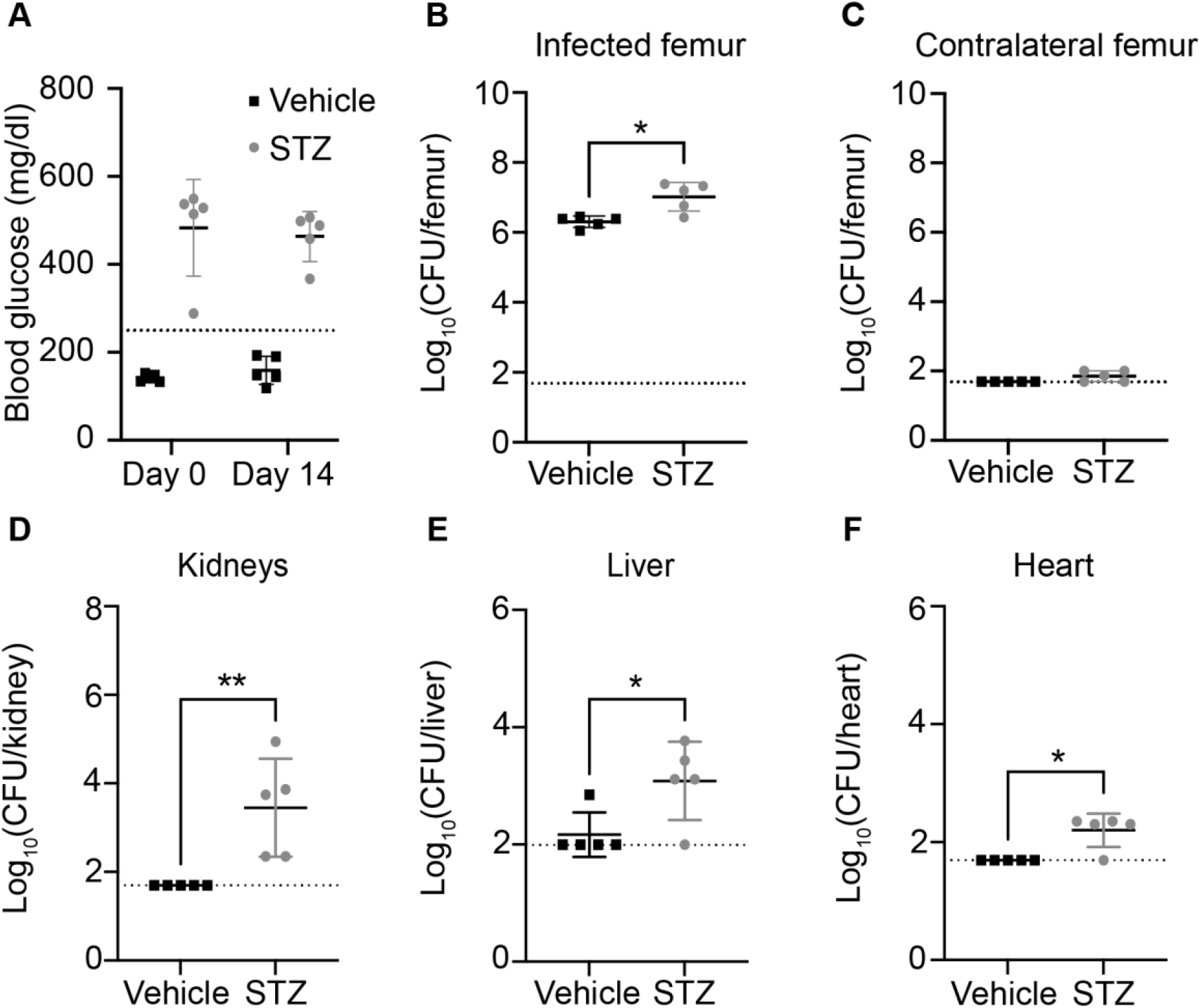
Chronic hyperglycemia increases *S. aureus* burdens and dissemination during osteomyelitis. Eight-week old male mice were treated with sodium citrate (vehicle) or streptozotocin (STZ) intraperitoneally for 5 days. 30 days after the final injection, mice were infected with 1×10^6^ CFU of WT *S. aureus* via intraosseous injection. (A) Blood glucose concentration was quantified from a tail vein bleed immediately prior to inoculation (day 0) and the day of sacrifice (day 14). Dotted line indicates hyperglycemia cut-off of 250 mg/dl. Mice were sacrificed at day 14 post-infection, and the bacterial burdens (CFU) were enumerated in (B) infected femur, (C) contralateral femur, (D) kidneys, (E) liver, and (F) heart. N = 5 mice per group. Dotted lines indicate limit of detection. Horizontal lines indicate mean, and error bars represent SD. Significance determined with Mann-Whitney test (B-F). *p<0.05, **p<0.01.

### Hyperglycemia increases bone loss during *S. aureus* osteomyelitis

To further characterize the pathogenesis of osteomyelitis during hyperglycemia, we measured bone loss following *S. aureus* infection. Significant bone damage and pathologic remodeling occur during *S. aureus* osteomyelitis in euglycemic animals.^18,23,25^ Due to the increased *S. aureus* burdens observed in hyperglycemic mice compared to euglycemic mice, we hypothesized that bone destruction would increase in the setting of hyperglycemia. STZ-treated hyperglycemic mice and vehicle-treated euglycemic mice were inoculated with 1×10^5^ CFU of *S. aureus* due to the increased infection severity with a 1×10^6^ CFU inoculum. We used microcomputed tomography (μCT) to quantify changes to bone structure in the *S. aureus*-infected femurs at 14 days post-infection. The infected femurs from hyperglycemic mice had greater cortical bone loss relative to the infected femurs from euglycemic vehicle-treated mice during acute hyperglycemia (**Fig. 3A**). Similar trends in cortical bone loss were observed in infected mice with chronic hyperglycemia (**Fig. S4A**). To further characterize the impact of acute and chronic hyperglycemia on bone homeostasis, we also measured changes in the trabecular bone volume. A decrease in trabecular bone volume relative to total volume (%BV/TV) was observed in the infected femurs from hyperglycemic mice compared to infected femurs from euglycemic vehicle-treated mice (**Fig. 3B**). Because chronic hyperglycemia has been shown to alter bone volume, we normalized the %BV/TV against the uninfected contralateral femur.^14,26^ Importantly, the %BV/TV in the hyperglycemic infected femurs normalized against the %BV/TV of the contralateral femurs remained significantly lower in the hyperglycemic mice compared to the euglycemic vehicle-treated mice, suggesting the trabecular bone loss in hyperglycemic mice is related to infection and not solely a function of baseline changes in bone volume (**Fig. 3C**). Trabecular bone thickness in infected hyperglycemic animals was lower than in the infected euglycemic vehicle-treated animals, with no differences observed in trabecular spacing or number (**Fig. 3D-F**). Similar trends in trabecular bone parameters were observed in the infected femurs of mice with chronic hyperglycemia (**Fig. S4B-F**). To further characterize tissue inflammation in hyperglycemic and euglycemic mice subjected to osteomyelitis, representative histological sections of *S. aureus*-infected femurs were stained with H&E. Hyperglycemic infected femurs exhibited greater signs of inflammation compared to the euglycemic vehicle-treated infected femurs (**Fig. S5**). Representative histological sections of infected femurs from chronic hyperglycemic mice exhibited similar pathology to acute hyperglycemic infected femurs (**Fig. S6**). Collectively, our data reveal that acute and chronic hyperglycemia contribute to greater pathologic bone destruction during *S. aureus* osteomyelitis compared to euglycemic infection. Due to the similar findings between acute and chronic hyperglycemic mice, we performed further experiments using the acute model of hyperglycemia.

**Figure 3.**
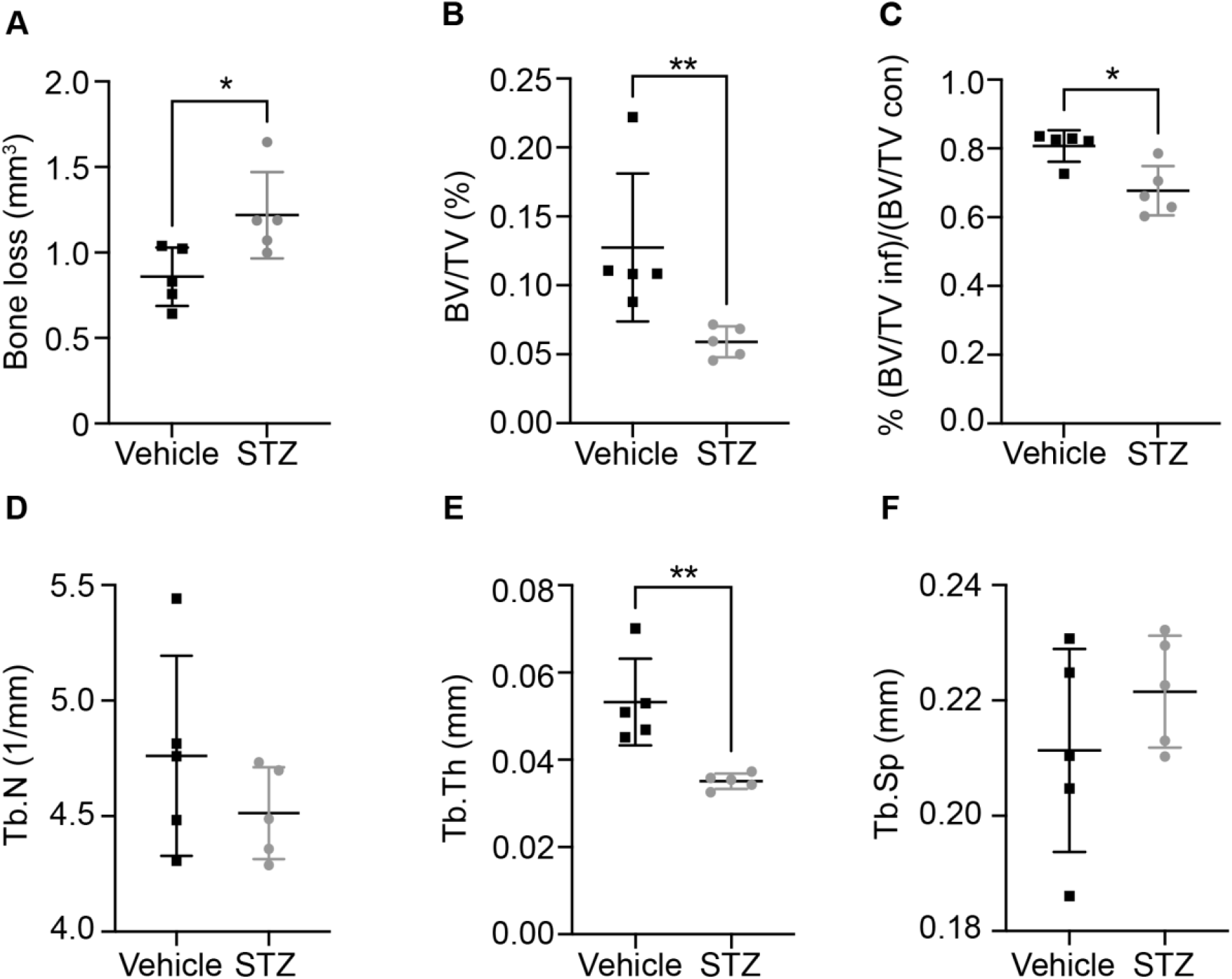
*S. aureus* incites greater bone destruction in acute hyperglycemic animals. Eight-week old male mice were treated with sodium citrate (vehicle) or streptozotocin (STZ) intraperitoneally for 5 days. 10 days after the final injection, mice were infected with 1×10^5^ CFU of WT *S. aureus* via intraosseous injection. At 14 days post-infection, the infected femur and contralateral femur were isolated for microcomputed tomography. (A) Cortical bone loss in infected femurs, (B) trabecular bone volume divided by total volume (BV/TV) of infected femurs, (C) BV/TV of infected femurs relative to contralateral femurs were quantified. (D) Trabecular number (Tb.N), (E) trabecular thickness (Tb.Th), and (F) trabecular spacing (Tb.Sp) were quantified in infected femurs. N = 5 mice per group. Horizontal lines indicate mean, and error bars represent SD. Significance determined with Mann-Whitney test. *p<0.05, **p<0.01.

### Genes contributing to the fitness of *S. aureus* in hyperglycemic animals

Hyperglycemia results in increased *S. aureus* burdens within infected femurs and greater dissemination to other organs in the context of osteomyelitis. To identify genes required for staphylococcal survival in hyperglycemic tissues, we performed TnSeq in hyperglycemic and euglycemic animals with a previously characterized USA300 LAC transposon insertion library.^19^ Groups of mice were treated with STZ or vehicle, and 10 days after the final treatment, mice were infected with 5×10^6^ CFU of the *S. aureus* transposon insertion library. An *in vitro* comparator consisted of growth in BHI for 24 hrs to identify genes that are essential for bacterial growth in a nutrient rich condition.

TnSeq identified 71 *S. aureus* genes as essential (defined as Dval<0.01) for survival during osteomyelitis in hyperglycemic mice, but not essential for growth *in vitro* or in euglycemic mice (**Fig. 4A** and **Table S1**). We also identified 61 *S. aureus* transposon mutants with compromised fitness (Dval>0.01 and <0.1) in hyperglycemic mice but not *in vitro* or in euglycemic mice (**Table S2**). Of these 132 genes, 45 encode hypothetical proteins, of which 28 genes are considered essential and 17 have compromised fitness. Of the remaining 87 genes, 61 have Kyoto Encyclopedia of Genes and Genomes (KEGG) identifiers. Of the 61 genes with KEGG identifiers, 24 are implicated in metabolic processes including glycolysis, glutamine metabolism, histidine metabolism, and purine/pyrimidine biosynthesis. Genes related directly or indirectly to glucose metabolism that were identified as either essential or compromised during osteomyelitis include and catabolite control protein A (*ccpA*, SAUSA300_1682), L-lactate dehydrogenase 1 (*ldh1*, SAUSA300_0235), dihydrolipoyl dehydrogenase (*lpdA*, SAUSA300_0996), and pyruvate ferredoxin oxidoreductase (SAUSA300_1182). Other metabolic genes identified as essential included carbamoyl phosphate synthase large subunit (*carB*, SAUSA300_1096), carbamoyl phosphate synthase small subunit (*carA*, SAUSA300_1095), and adenine phosphoribosyltransferase (*apt*, SAUSA300_1591).

**Figure 4.**
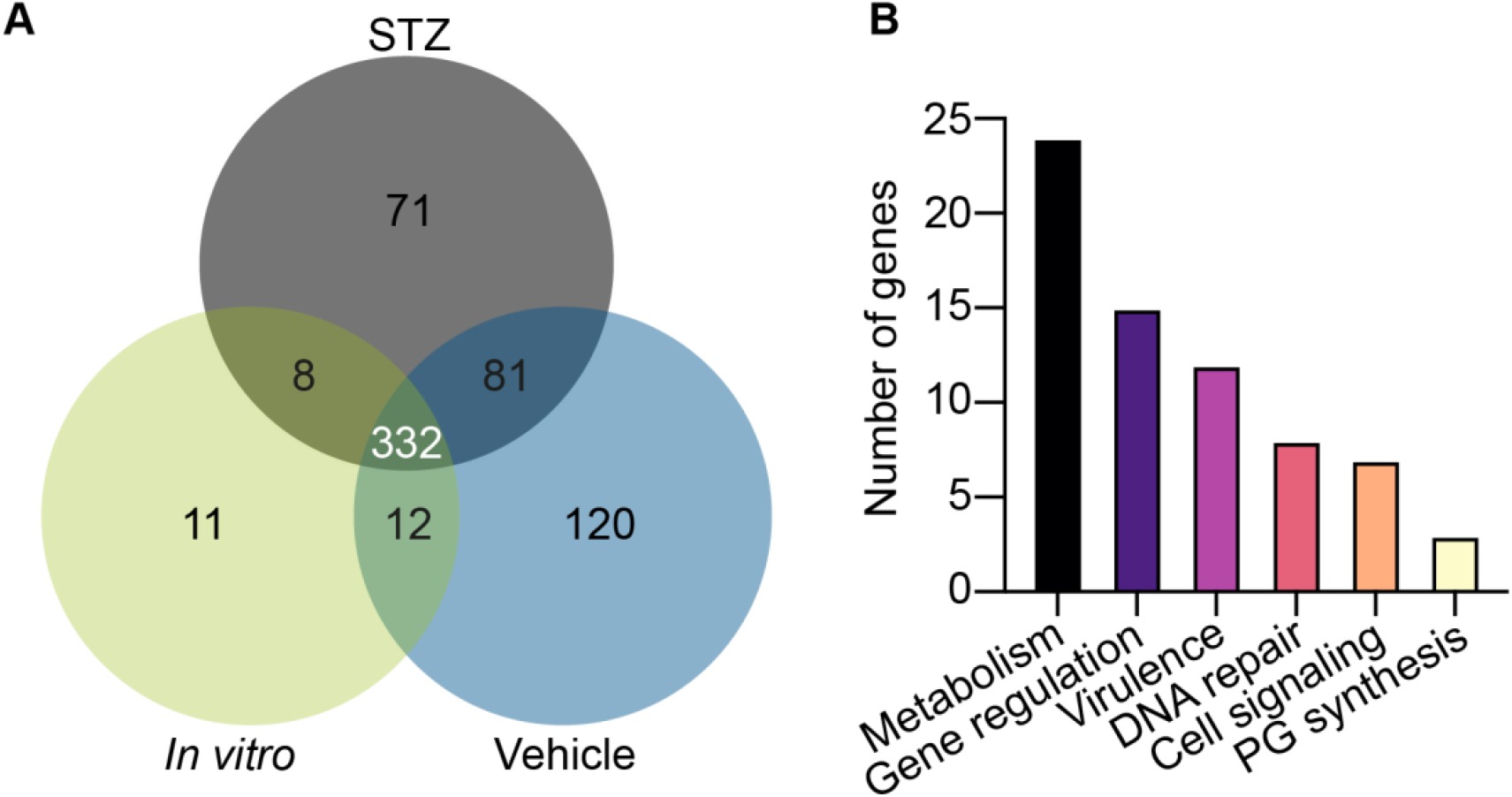
Transposon sequencing reveals genes essential for *S. aureus* survival during osteomyelitis in mice with hyperglycemia. Eight-week old male mice were treated with sodium citrate (vehicle) or streptozotocin (STZ) intraperitoneally for 5 days. 10 days after the final injection, mice were infected with 5×10^6^ CFU of the *S. aureus* TnSeq library via intraosseous injection. At day 4 post-infection, bacteria were recovered and Illumina Sequencing was used to identify the abundance of *S. aureus* mutants in each condition. (A) Based on Dval calculations, the number of essential genes (Dval<0.01) *in vitro* in BHI, *in vivo* in STZ-treated hyperglycemic mice, and *in vivo* in euglycemic vehicle-treated mice were enumerated. (B) The annotated genes essential for *S. aureus* survival (Dval <0.01) and the mutants with compromised fitness (Dval>0.01 and <0.1) only in the condition of osteomyelitis during hyperglycemia were organized into categories of metabolism, gene regulation, virulence, DNA repair, cell signaling, and peptidoglycan (PG) synthesis. N = 12 mice per *in vivo* group, pooled in samples of 2 for n = 6 in sequencing analysis.

In addition to metabolic pathways identified as important for *S. aureus* growth in osteomyelitis during hyperglycemia, the remaining genes identified by TnSeq are broadly involved in bacterial stress responses. These genes can be classified by roles in DNA repair, cell signaling, gene regulation, peptidoglycan synthesis, and virulence (**Fig. 4B**). One virulence gene identified as uniquely essential for *S. aureus* growth during osteomyelitis in hyperglycemic mice was *sodA* (SAUSA300_1513), which encodes superoxide dismutase A. SodA is responsible for detoxifying ROS via conversion of superoxide radicals to hydrogen peroxide.^27^ Due to the hyperinflammation during hyperglycemia and presence of greater ROS concentrations, we sought to understand how SodA facilitates *S. aureus* growth in the presence of elevated glucose.

### SodA facilitates *S. aureus* survival in high glucose *in vitro*

While host cells produce ROS during inflammation, *S. aureus* can also produce ROS intrinsically in response to glucose metabolism that results in an electron transport chain bottleneck, catalyzing reduction of oxygen.^28^ To combat ROS, the *S. aureus* genome encodes two superoxide dismutases (SODs), *sodA* and *sodM*^27,29^ The *sodA* gene was identified as essential for growth during osteomyelitis in hyperglycemic mice, while *sodM* was not essential for growth *in vitro*, in euglycemic vehicle-treated osteomyelitis infection, or in osteomyelitis in hyperglycemic mice. To evaluate the roles of each SOD in *S. aureus* survival in elevated glucose in the absence of other host stressors, we examined survival of WT, *sodA* mutant, and *sodM* mutant strains over 5 days in TSB (250 mg/dl glucose) or in TSB with an additional 500 mg/dl of glucose by quantifying CFU every 24 hrs.^28^ While WT and the *sodM* mutant survive in TSB and TSB with glucose to a similar extent over the course of 5 days, the *sodA* mutant had a growth defect in TSB with glucose compared to WT by day 2 (**Fig. 5A**). Prior studies suggests that oxygen is critical for SOD activity, which led us to hypothesize that differences in growth between WT and the SOD mutants would be minimized under conditions of limited oxygen.^27,30^ In keeping with this hypothesis, WT *S. aureus* grew similarly to the *sodA* mutant strain under microaerobic conditions (**Fig. 5B**). Although *S. aureus* cultures with high glucose became more acidic over time, changes in growth between *S. aureus* strains were not explained by differences in pH (**Fig. 5C).** Complementation with *sodA in cis* rescues the growth the *sodA* mutant *in vitro* when grown aerobically in TSB with glucose (**Fig. 5D**). Consistent with prior studies, these findings support a role for SodA in detoxifying intrinsically generated superoxide.^31,32^

**Figure 5.**
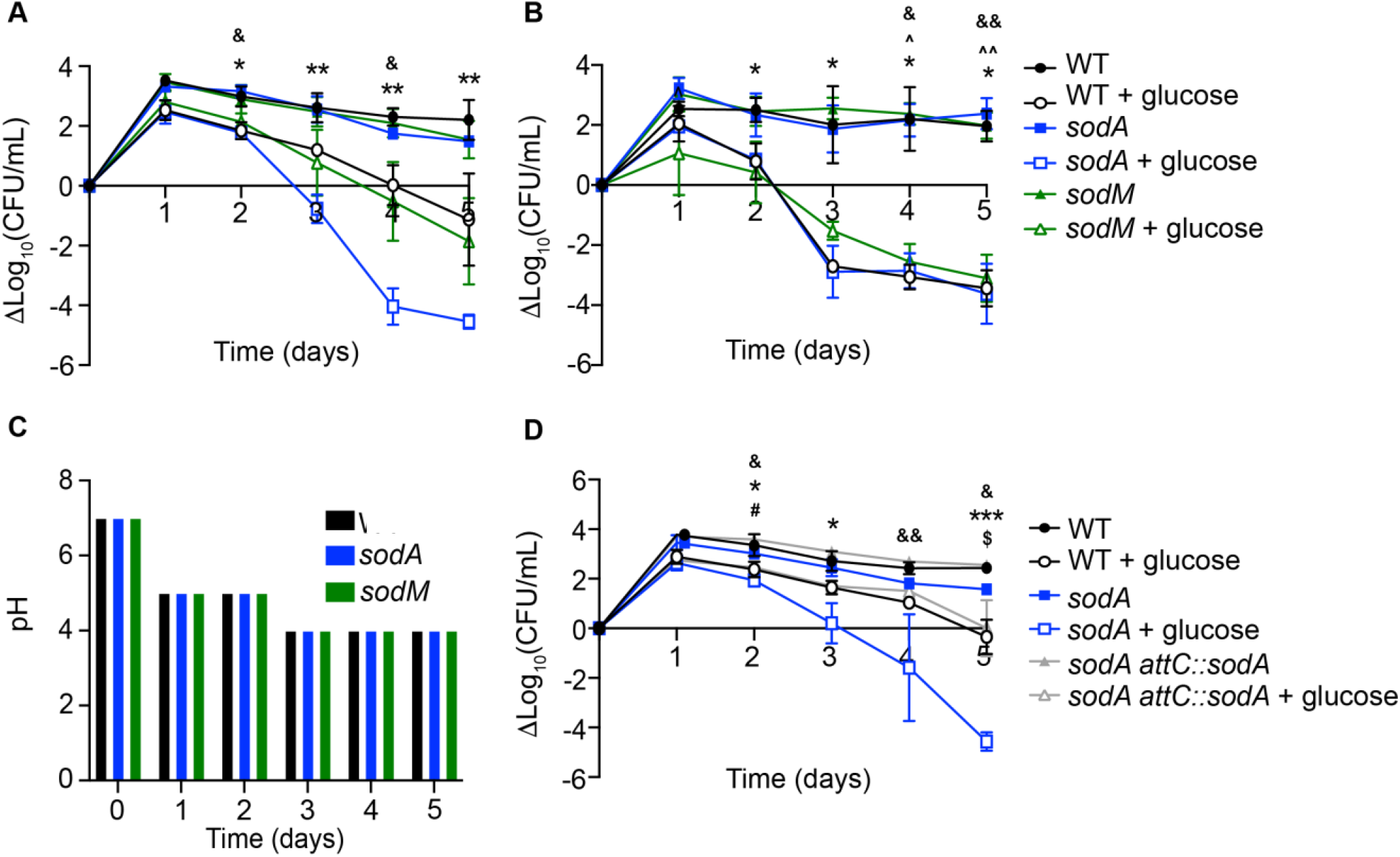
*S. aureus sodA* is required for survival during culture in high glucose. WT, *sodA::tet, sodM::erm*, and *sodA::tet sodM::erm S. aureus* strains were grown in 10 mL TSB with (“+ glucose”) and without 500 mg/dl glucose in flasks covered with (A) foil (aerobic) or (B) capped (microaerobic) shaking at 37°C. CFU were quantified every 24 hrs over the course of 5 days and normalized to time 0 hr. (C) pH was measured every 24 hrs over the course of 5 days in TSB with 500 mg/dl glucose grown in aerobic microaerobic conditions. (D) WT, *sodA*::tet, and *sodA::tet attC::sodA S. aureus* were grown in 10 mL TSB and TSB with 500 mg/dl glucose (“+ glucose”) in flasks covered with foil (aerobic). N = 2 technical replicates, and n = 3 biological replicates. Line represents mean (A, B, and D). Significance determined with two-way ANOVA and Dunnett’s multiple comparisons test. *p<0.05, **p<0.01, ***p<0.001 *sodA*::tet TSB + glucose; &p<0.05, &&p<0.01 WT TSB + glucose; ^p<0.05, ^^p<0.01 *sodM*::erm TSB + glucose; # *sodA*::tet *attC::sodA* TSB + glucose; $p<0.01 *sodA*::tet TSB. All comparisons made to WT in TSB.

### SodA enhances *S. aureus* survival during osteomyelitis in hyperglycemic mice

To validate the role of *S. aureus* SODs *in vivo*, we assessed the CFU burdens of WT, *sodA* mutant, and *sodM* mutant *S. aureus* in osteomyelitis mono-infections during hyperglycemia. Following STZ treatment, we inoculated mice with 1×10^5^ CFU of each mutant strain or WT. Over the course of the infection, mice were monitored for changes in weight. Mice infected with *sodA* and *sodM* mutants had significantly less weight loss at multiple days post-infection compared to mice infected with WT (**Fig. 6A**). *S. aureus* burdens were lower in femurs of mice infected with a *sodA* mutant compared to WT at day 14 post-infection (**Fig. 6B**). Furthermore, while the kidneys and liver from hyperglycemic mice infected with the SOD mutant strains did not exhibit differences in bacterial burdens compared to WT, the hearts of animals infected with the *sodA* mutant had lower *S. aureus* burdens compared to WT infected mice (**Fig. 6C-E**). These data confirm the importance of SodA in promoting *S. aureus* osteomyelitis in the setting of hyperglycemia.

**Figure 6.**
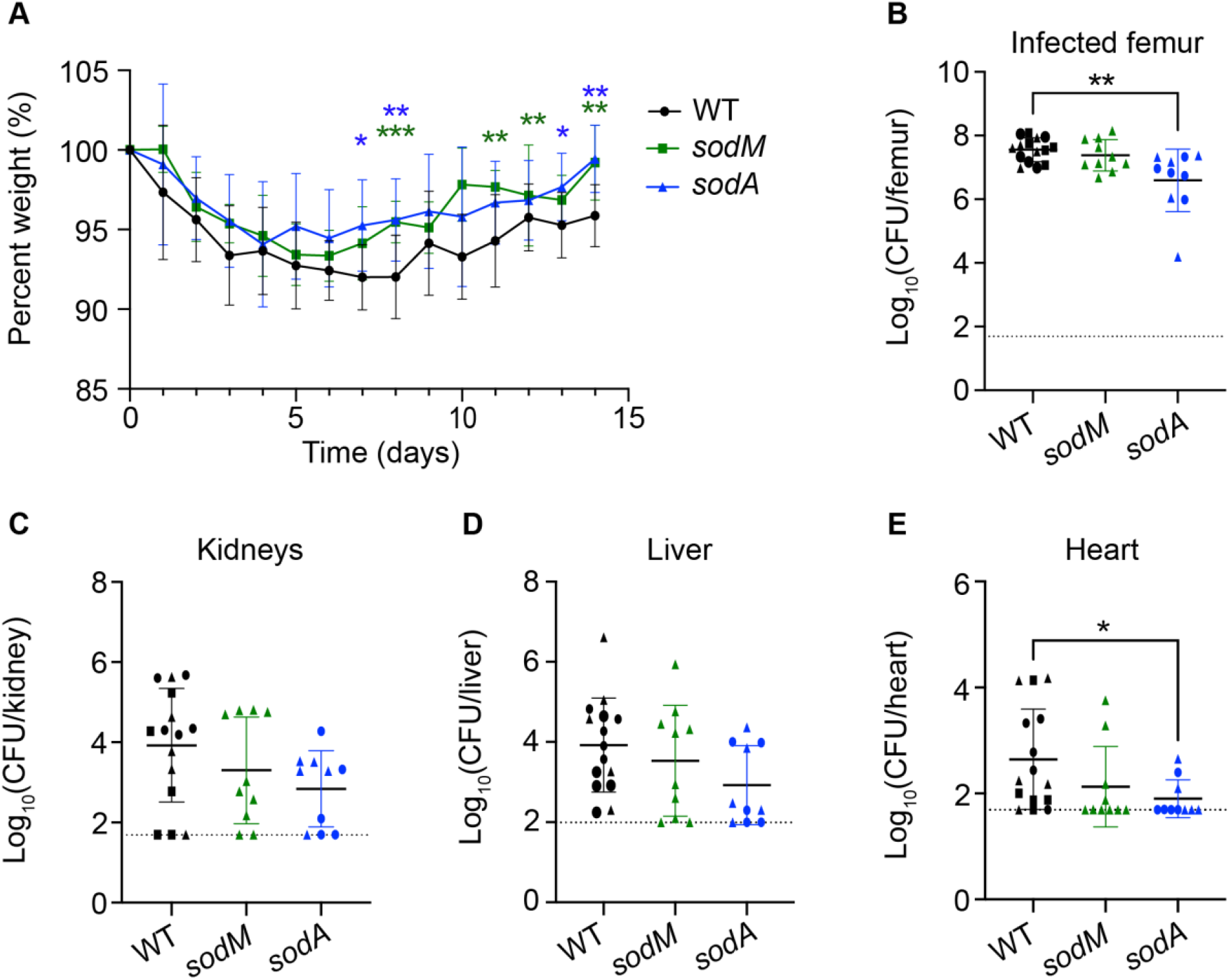
SodA is important for *S. aureus* survival during osteomyelitis in hyperglycemic mice. Eight-week old male mice were treated with streptozotocin (STZ) intraperitoneally for 5 days. 10 days after the final injection, mice were infected with 1×10^5^ CFU of WT, *sodA::tet*, and *sodM::erm S. aureus* via intraosseous injection. (A) Weights were recorded every 24 hrs and normalized to the starting weight of each animal on the day of infection (percent weight). Mice were sacrificed at day 14 post-infection, and the bacterial burdens (CFU) were enumerated in (B) infected femur, (C) kidneys, (D) liver, and (E) heart. Different shapes indicate data collected from distinct experiments. WT n = 15 mice, all other strains n = 10 mice per group. Dotted lines indicate limit of detection. Horizontal lines indicate mean, and error bars represent SD. Significance determined with two-way ANOVA and Dunnett’s multiple comparisons test (A) and Kruskal-Wallis with Dunn’s multiple comparisons test (B-E). *p<0.05, **p<0.01, ***p<0.001. All comparisons made to WT.

To assess whether *S. aureus* SODs are essential in osteomyelitis in euglycemic mice, we inoculated vehicle-treated control mice with 1×10^5^ CFU WT, *sodA* mutant, or *sodM* mutant *S. aureus* and measured bacterial burdens in the femur 14 days post-infection. Neither of the SOD mutant infected mice showed differences in percent starting weight compared to WT (**Fig. 7A**). WT, *sodA* mutant, and *sodM* mutant infected animals showed no differences in burdens in the infected femurs at day 14 post-infection (**Fig. 7B**). These data suggest that the fitness defect of the *sodA* mutant is unique to hyperglycemic osteomyelitis.

**Figure 7.**
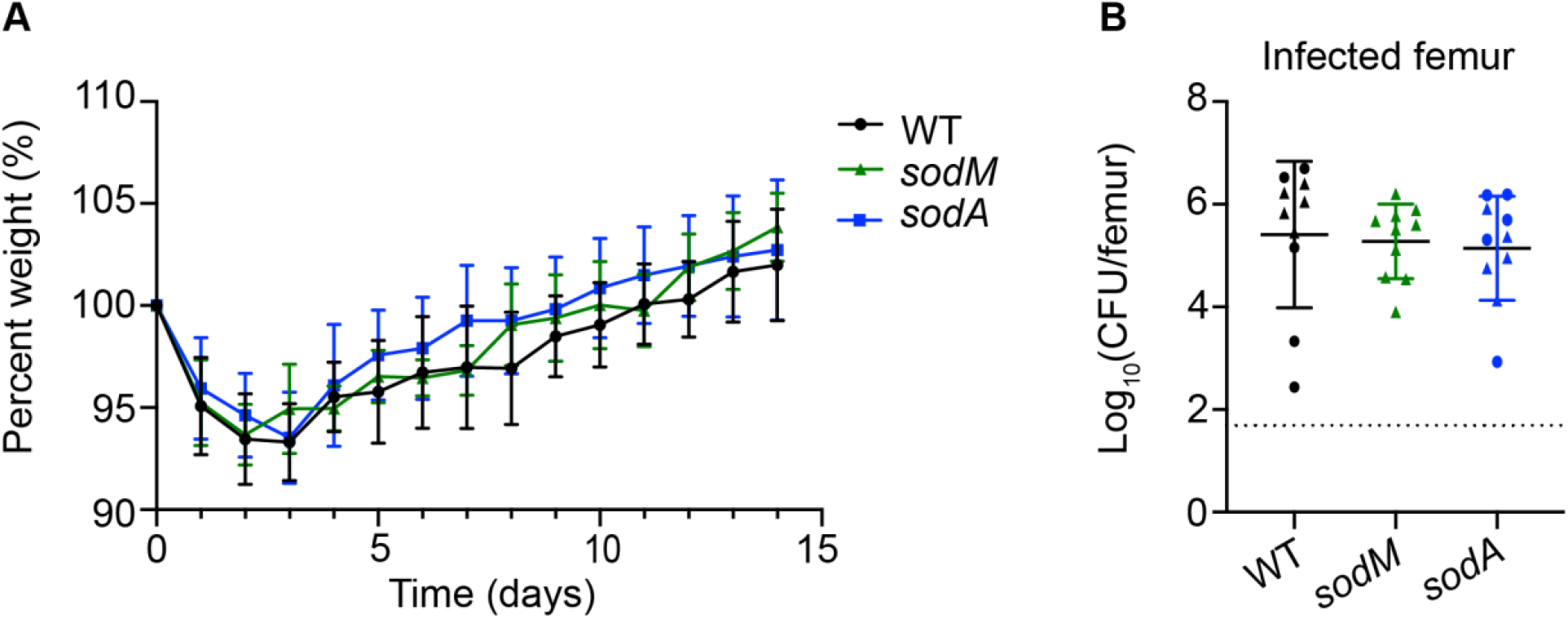
SodA is not necessary for *S. aureus* survival during osteomyelitis in euglycemic mice. Eight-week old male mice were treated with sodium citrate (vehicle) intraperitoneally for 5 days. 10 days after the final injection, mice were infected with 1×10^5^ CFU of WT, *sodA::tet*, and *sodM::erm S. aureus* via intraosseous injection. (A) Weights were recorded every 24 hrs and normalized to the starting weight of each animal on the day of infection (percent weight). (B) Mice were sacrificed at day 14 post-infection, and the bacterial burdens (CFU) were enumerated in infected femurs. Different shapes indicate data collected from distinct experiments. N = 10 mice per group. Dotted line indicates limit of detection. Horizontal lines indicate mean, and error bars represent SD. Significance determined with two-way ANOVA and Dunnett’s multiple comparisons test (A) and Kruskal-Wallis with Dunn’s multiple comparisons test (B). All comparisons made relative to WT.

## DISCUSSION

Using a chemically-induced model of hyperglycemia, we discovered increased *S. aureus* burdens and significantly greater bone loss during hyperglycemic infection compared to euglycemic infection in a post-traumatic osteomyelitis model. We also discovered increased dissemination of *S. aureus* to other organs in hyperglycemic mice. The gene encoding SodA was found to be critical for *S. aureus* survival during hyperglycemic infection and for *S. aureus* growth *in vitro* in the presence of high glucose. These findings are consistent with prior clinical studies that correlated poor *S. aureus* infection outcomes in hyperglycemic individuals, both with and without diabetes.^33,34^ This work further reveals a *S. aureus* virulence factor that contributes to increased osteomyelitis pathogenesis during hyperglycemia and supports the hypothesis that metabolic comorbidities shape the essential *S. aureus* genes required for invasive infection.

Multiple murine *S. aureus* infection models have identified increases in infection severity in the setting of hyperglycemia, including footpad infections in STZ-induced hyperglycemic mice.^1,35–37^ More severe infections were also observed in STZ-induced hyperglycemic mice in an implant-related orthopedic infection model, as measured by *S. aureus* burdens, bone density, and biofilm formation.^38^ Models of type 2 diabetes likewise revealed increased osteomyelitis infection severity associated with hyperglycemia.^39–41^ Elevated *S. aureus* burdens were observed in skin infection due to aberrant neutrophil chemotaxis, dysregulated abscess formation, and impaired wound healing during hyperglycemia.^42,43^ Our study contributes to a greater understanding of how hyperglycemia influences *S. aureus* infection pathogenesis by revealing changes in bacterial survival during post-traumatic osteomyelitis as well as significantly greater infection-induced bone loss during hyperglycemia.

In a prior study, we identified genes involved in glucose metabolism and virulence as essential for *S. aureus* survival and disease pathogenesis *in vivo* during osteomyelitis.^16,18,23^ To identify genes that contribute to increased pathogenesis of *S. aureus* osteomyelitis during hyperglycemia, we conducted TnSeq, comparing *S. aureus* survival in hyperglycemic mice to euglycemic mice. Similar to our prior TnSeq study, genes involved in purine/pyrimidine and amino acid metabolism were identified as essential for *S. aureus* growth during osteomyelitis in hyperglycemic mice, including *carA*, *carB*, and *aptA*. Both *carA* and *aptA* are important for *S. aureus* survival in nitric oxide stress.^19^ Furthermore, *carA* is critical for *S. aureus* extracellular survival in the presence of high H2O2.^44^ Purine and pyrimidine synthesis enable *S. aureus* to persist in the presence of hydroxyl radicals that oxidize base and ribose moieties of DNA, creating lesions that require repair.^45^ The gene *ccpA* was also uniquely essential for *S. aureus* growth during osteomyelitis in hyperglycemic mice, linking the abundance of glucose in tissues with transcriptional control of genes involved in gluconeogenesis, the TCA cycle, and cell adherence and immune evasion.^46–49^

TnSeq also revealed the importance of *sodA* for *S. aureus* survival during osteomyelitis in hyperglycemic mice. SodA has been implicated in *S. aureus* stress responses during aerobic metabolism and in the presence of ROS produced by innate immune cells.^27,32^ Additionally, *sodA* was previously identified as essential for *S. aureus* growth in skin infections in STZ-induced hyperglycemic mice.^36^ SodA catalyzes the detoxification of superoxide into hydrogen peroxide, which is further broken down by catalase, and both SOD and catalase activity of bacteria have been correlated to virulence.^27,50–52^ Production of superoxide is a known innate immune defense mechanism used to kill intracellular phagocytosed bacteria.^53,54^ Previous publications revealed that neutrophils have a decreased capacity for respiratory burst and limited ROS production during hyperglycemic infection.^5,43,55,56^ However, oxidative stress from altered metabolism of endothelial cells increases the amount of ROS in tissues of hyperglycemic patients.^6,57,58^ Superoxide can also be produced intrinsically by *S. aureus* in response to carbon overflow metabolism. Carbon overflow occurs when *S. aureus* breaks down glucose through glycolysis and results in acetate accumulation.^28^ The accumulation of acetate and NADH leads to a bottleneck in the electron transport chain that catalyzes reduction of oxygen, producing ROS.^59^ Based on these findings, we hypothesized that a *sodA* mutant would have a defect in survival in high glucose. We observed decreased *S. aureus* growth in glucose-rich media over 5 days in the absence of a functional *sodA* gene.^60^ These data are consistent with a model of hyperglycemic skin infection whereby dysfunctional phagocytes were observed to consume less glucose during hyperglycemia, thereby eliminating *S. aureus* competition for glucose and potentiating S. *aureus* virulence.^35^ These data suggest that the defect in survival of a *sodA* mutant may be related to intrinsic *S. aureus* ROS production during hyperglycemia.

The *S. aureus* genome encodes two SOD genes, *sodA* and *sodM*.^27,29^ The SODs are critical for *S. aureus* survival in the presence of superoxide during distinct phases of growth, with SodA functioning to detoxify ROS during exponential phase, while SodM has a greater role during stationary phase.^29^ Both *S. aureus* SODs can utilize manganese as a co-factor, while SodM can also use iron.^31^ The distinct characteristics of SodA and SodM may facilitate *S. aureus* survival in the presence of ROS in distinct tissues and/or nutritional microenvironments. Because SodA activity may be compensated by SodM, we decided to interrogate the roles of both genes in osteomyelitis during hyperglycemia. Mutating *sodA* decreases *S. aureus* survival in hyperglycemic infection, while mutating *sodM* does not affect *S. aureus* survival, further supporting the importance of SodA in this model of infection. In comparison, prior studies in euglycemic mice found that both *sodA* and *sodM* are essential for full *S. aureus* virulence following intravenous infection.^61^ These data suggest that there could be tissue and microenvironment-specific factors that influence the need for *S. aureus* SODs to support bacterial survival *in vivo*. We did not observe differences in the ability of WT and SOD mutants to survive osteomyelitis in euglycemic vehicle-treated mice. These findings are consistent with a skin abscess model of infection where there was no difference in abscess formation with a *sodA* mutant strain compared to WT in euglycemic mice.^27^

There are some limitations to this study that should be considered. We chose to induce hyperglycemia with STZ to model increased blood glucose while minimizing confounding physiological changes associated with obesity, age, or the need for specialized housing.^62^ Limitations of this model include that STZ can be toxic to other organs, and the conclusions may not be generalizable to other models of hyperglycemia.^62,63^ To model osteomyelitis, we used an established post-traumatic model, inoculating *S. aureus* directly into the femurs.^18^ This model does not effectively reproduce the characteristic clinical progression of contiguous wound dissemination, which is commonly observed with diabetic foot wounds. Finally, TnSeq has inherent limitations, including the ability of compensatory mutants to rescue the growth of transposon mutants via nutrient sharing or other community interactions.

Future studies should include other models of osteomyelitis, such as foot pad infections, to observe whether osteomyelitis infection dynamics are recapitulated in contiguous wound infection models. Additionally, studies are needed to assess which host processes influence the role of SodA in infection. Ablating the function of neutrophils in healthy mice prior to infecting with a *S. aureus sodA* mutant may reveal the contributions of innate immune dysregulation versus the contribution of circulating blood glucose concentration. Additionally, *S. aureus* glucose transporter mutants can be used to assess the ability of *S. aureus* to survive in hyperglycemic conditions when it does not have the ability to use exogenous glucose.^64^ Measuring changes in metabolites and ROS within the microenvironment of infected femurs in hyperglycemic mice is also an important future direction.

Taken together, the data in this manuscript reveal that both acute and chronic hyperglycemia increase *S. aureus* infectious burden, dissemination, and bone destruction during osteomyelitis. We identified 71 genes that are uniquely essential for *S. aureus* growth during osteomyelitis in hyperglycemic mice compared to euglycemic osteomyelitis. Of these genes, *sodA* was further studied due to its role in detoxifying ROS, a biproduct of glucose carbon overflow metabolism. The results of this study highlight a bacterial virulence gene that contributes to exacerbated infection during hyperglycemia and provide a strong rationale for continued investigation into mechanisms of enhanced musculoskeletal disease in the context of hyperglycemia.

## METHODS

### Bacterial strains and culture conditions

Unless otherwise stated, experiments were performed with *S. aureus* USA300 lineage strain AH1263, an erythromycin and tetracycline sensitive derivative of strain LAC, which served as wildtype (WT). Strain *sodA*::tet and *sodM::erm* in the AH1263 background were created via phi-85-mediated phage transduction of *sodA*::tet and *sodM*::erm from the Newman background.^29,31^ The *sodA* complementation construct was created by amplifying *sodA* and its endogenous promoter with primer sequences CTAGCTCTAGATGAGATTTATGCACATTTGGTCA and CTAGCGGTACCTTTATTTTGTTGCATTATATAATTCG. The *sodA* sequence was ligated into pJC1111 and integrated into the chromosome at attachment (*attC*) sites, as previously described.^65^ All bacterial cultures were grown overnight in 5 ml Tryptic Soy Broth (TSB) at 37°C with shaking at 180 rpm, except as otherwise noted. 10 μg/ml erythromycin, 2 μg/ml tetracycline, or 0.1 mM cadmium chloride were added to cultures with strains possessing antibiotic resistance markers.

### Murine model of osteomyelitis

All animal experiments were reviewed and approved by the Institutional Animal Care and Use Committee at Vanderbilt University Medical Center and performed in accordance with NIH guidelines, the Animal Welfare Act, and US Federal law. Six - eight week male C57Bl/6J mice were obtained from Jackson Laboratories (stock #000664) and intraperitoneally injected daily with 200 μL of 0.1 mM sodium citrate (vehicle) or 40-60 mg streptozotocin (STZ) in 200 μL of 0.1 mM sodium citrate for 5 days to induce hyperglycemia. Acute hyperglycemia infections were performed 10 days after the final intraperitoneal STZ or sodium citrate injection, and chronic hyperglycemia infections were performed 30 days the final intraperitoneal STZ or sodium citrate injection. Blood glucose concentration was quantified from a tail vein bleed immediately prior to inoculation (day 0) and on the day of sacrifice (day 14). STZ-treated mice below the hyperglycemic threshold of 250 mg/dl were removed from the study, except as otherwise noted. Osteomyelitis was induced with ~1 x 10^6^ or ~1 x 10^5^ CFU in 2 μl via intraosseous injection into the femurs, as previously reported.^18^ Mice were weighed daily and monitored for disease progression. 14 days post-infection, mice were sacrificed, and the infected femurs were either homogenized for CFU enumeration or fixed for micro-computed tomography (μCT) (see below). For CFU enumeration, infected femur, contralateral femur, kidneys, liver, and heart were homogenized in Cell Lytic buffer (Sigma) and plated on tryptic soy agar (TSA). Limits of detection based on volume of homogenate plated were 49 CFU per femur, heart, and kidneys and 99 CFU per liver.

### Micro-computed tomography analysis of femurs

Infected and contralateral femurs were harvested from mice at day 14 post-infection. Femurs were fixed in 10% neutral buffered formalin for 2 days then moved to 70% ethanol and stored at 4°C. Fixed femurs were scanned with a μCT50 Scanco instrument (Scanco Medical, Switzerland) and analyzed with μCT Tomography V6.3-4 software (Scanco USA, Inc.). The diaphysis and distal epiphysis of each femur were imaged with 10.0 μm voxel size at 70 kV, 200 μA with an integration time of 350 ms in 10.24 mm view. 1088 slices were obtained to include the diaphysis surrounding the cortical defect formed during inoculation as well as trabecular bone in the distal femur. Three-dimensional reconstructions were analyzed to quantify cortical bone destruction surrounding the inoculation site (mm^3^). Trabecular bone volume per total volume (%), trabecular number (1/mm), trabecular thickness (mm), and trabecular spacing (mm) were quantified within the epiphyses, as previously described.^18,25^

### Bone histology

After μCT imaging, femurs were decalcified in 20% EDTA for 4 days at 4°C. Decalcified femurs were processed into paraffin, embedded, and sectioned through the infectious nidus and bone marrow cavity at 4 μm thickness with a Leica RM2255 microtome. Sections were stained with hematoxylin and eosin (H&E). A Leica SCN400 Slide Scanner was used to scan stained femur sections at 20X, and images were uploaded to and analyzed within the Digital Slide Archive (Vanderbilt University Medical Center).

### Transposon sequencing analysis of experimental osteomyelitis

USA300 LAC transposon library aliquots were obtained and expanded in 10 ml BHI in 50 ml Erlenmeyer flasks loosely covered with foil for 6 hrs at 37°C with shaking at 180 rpm.^19,66^ The expanded library was collected by centrifugation, aliquoted for individual experiments, and thawed on ice as needed, as previously described.^66^ Briefly, library aliquots were centrifuged at 200 x g for 8 min at 4°C and resuspended in cold, sterile PBS to achieve an inoculum concentration of ~2.5 x 10^9^ CFU/ml. 2 μl of inoculum was delivered (final concentration of ~5 x 10^6^ CFU) via intraosseous injection into the femurs of C57Bl/6J male mice treated with vehicle or STZ, as described above. Mice were sacrificed at day 4 based on prior studies.^23^ Femurs were homogenized in 500 μl of cold, sterile PBS. 150 μl of bone homogenate from two mice were pooled in 4.7 ml BHI in 50 ml Erlenmeyer flasks loosely covered with foil for a 2 hr outgrowth step at 37°C with shaking at 180 rpm. Following outgrowth, host debris was allowed to settle to the bottom of the culture, and the top fraction was transferred to a new conical on ice. The top fraction was pelleted at 8000 x g for 8 min at 4°C, resuspended in an equal volume of 20% glycerol BHI, and stored at - 80°C. In parallel, 2 μl of prepared inoculum was inoculated into 50 ml of BHI in a 250 ml Erlenmeyer flask to serve as an *in vitro* comparator. After 24 hrs of growth at 37°C with shaking at 180 rpm and covered loosely with foil, the cultures were pelleted at 8000 x g for 8 min at 4°C, supernatant was discarded, and samples were stored at −80°C.

### Library preparation and analysis of transposon sequencing

Genomic DNA was isolated with a phenol:chloroform:isoamyl alcohol protocol as described previously.^66^ The DNA was then sheared to ~350 bp by sonication using a Covaris LE220 instrument. Libraries were prepared for sequencing with the homopolymer tail-mediated ligation PCR technique.^67^ Terminal deoxytransferase was used to generate a poly(C)-tailed sequence on the 3’ end of the DNA fragments. The transposon junctions were amplified with two rounds of nested PCR and multiplexed with 8 bp indexing primers. The indexed DNA fragments were sequenced on an Illumina Hi-Seq 2500 (Tufts University Core Facility). Reads were trimmed, filtered for quality, and mapped to *S. aureus* FPR3757 accession number NC007793. A Dval score was assigned to each gene based on the number of reads within a given gene in a sample divided by the predicted number of reads for the gene considering its size and total sequencing reads for the given sample. Genes with a Dval between 0.1 and 0.01 were considered compromised in each condition, and genes with a Dval of ≤0.01 were considered essential.

### Comparative *S. aureus* growth analysis *in vitro* in different concentrations of glucose

Overnight cultures of WT and mutant strains were washed in PBS and back diluted 1:1000 into 10 ml TSB with and without 500 mg/dl added glucose in 50 ml Erlenmeyer flasks. Cultures were grown at 37°C with shaking at 180 rpm either loosely covered with foil (aerobic) or plugged with a rubber stopper (microaerobic). Viable CFU were measured by serial diluting cultures and plating on TSA at the indicated timepoints. Growth was reported as Log_10_CFU/ml compared to CFU enumerated at 0 hrs. pH was measured over the course of the experiment with pH test strips (FisherScientific 13-640-516).

### Graphical and Statistical Analysis

Statistical analyses were performed with Prism 9.0 (GraphPad Software). Data were checked for normality prior to statistical analysis. In comparisons of two groups, including the comparisons made for CFU burdens and μCT parameters, Mann-Whitney tests were used. To assess the importance of SOD genes for *S. aureus* survival over time *in vitro*, two-way ANOVAs with post hoc Dunnett’s multiple comparisons tests were used to compare the influence of the genotype at each time point. Two-way ANOVAs with post hoc Dunnett’s multiple comparisons tests were also used to compare the percent weight of animals infected with different *S. aureus* strains or mouse treatment (STZ- or vehicle-treated) over 14 days. To measure changes in *S. aureus* survival *in vivo* in experiments with 3 or more bacterial strains, one-way ANOVAs with Kruskal-Wallis and Dunn’s multiple comparisons were used due to the non-Gaussian distribution of the data. Changes in *S. aureus* bacterial burden at day 14 in vehicle-treated, STZ-treated not hyperglycemic, and STZ-treated hyperglycemic mice were compared with a one-way ANOVA and Tukey’s multiple comparisons test.

## ACKNOWLEDGMENTS

JEC was supported by the National Institute of Allergy and Infectious Diseases (NIAID) grants R01AI145992, R01AI161022, and R01AI132560 and by a Burroughs Wellcome Fund Career Award for Medical Scientists. CEB was supported by the NIAID Chemical Biology of Infectious Diseases T32, T32AI112541. NBK was funded by T32AI138932. ERG was supported by T32HL094296. JRJ was supported by funds from the Department of Medicine, Vanderbilt University Medical Center. SHP was supported by the Department of Veterans Affairs grant 1IK2BX005401 and by funds from the Department of Medicine, Vanderbilt University Medical Center. CBI was funded by the NIAID K22AI155927. CHS was supported by R01HL124159-01, DK122147-01A1, and AI149207A. Research support for the μCT50 and computer cluster was provided by National Institutes of Health (https://www.nih.gov) grant S10RR027631.

We would like to acknowledge Sasi Uppuganti and the Vanderbilt University Institute of Imaging Sciences (VUIIS) for assistance with μCT imaging and Dr. Eric Skaar (Vanderbilt University) for providing Newman *sodM*::erm and *sodA*::tet strains. We thank Dr. Anthony Richardson (University of Pittsburg) for providing the TnSeq library and the Tufts University Core Facility – Genomics Core for sequencing the transposon library samples. We also thank Dr. Erin Green (Vanderbilt University), Dr. Albert Tai (Tufts University), and Gloria Lam (Tufts University) for assisting with the TnSeq data analysis. Finally, we would like to acknowledge the Cassat laboratory members for providing feedback on data throughout the development of the project and for proofreading the manuscript.

